# *In-vitro* resensitization of multidrug resistant clinical isolates of *Enterococcus faecium* and *faecalis* through phage-antibiotic synergy

**DOI:** 10.1101/2024.06.17.599392

**Authors:** Pooja Ghatbale, Govind Prasad Sah, Sage Dunham, Ethan Khong, Alisha Blanc, Alisha Monsibais, Andrew Garcia, Robert T. Schooley, Ana G. Cobián Güemes, Katrine Whiteson, David T. Pride

## Abstract

Bacteriophages are an increasingly attractive option for the treatment of antibiotic resistant infections, but their efficacy is difficult to discern due to confounding effects of antibiotics. Phages are generally delivered in conjunction with antibiotics, and thus, when patients improve it’s unclear whether the phages, antibiotics or both are responsible. This question is particularly relevant for enterococcus infections, as limited data suggest phages might restore antibiotic efficacy against resistant strains. Enterococci can develop high-level resistance to vancomycin, a primary treatment. We assessed clinical and laboratory isolates of *Enterococcus faecium* and *Enterococcus faecalis* to determine whether we could observe synergistic interactions between phages and antibiotics. We identified synergy between multiple phages and antibiotics including linezolid, ampicillin, and vancomycin. Notably, antibiotic susceptibility did not predict synergistic interactions with phages. Vancomycin resistant isolates (n=6) were eradicated by the vancomycin-phage combination as effectively as vancomycin susceptible isolates (n=2). Transcriptome analysis revealed significant gene expression changes under antibiotic-phage conditions, especially for linezolid and vancomycin, with upregulated genes involved in nucleotide and protein biosynthesis and downregulated stress response and prophage-related genes. While our results do not conclusively determine the etiology of the observed synergistic interactions between antibiotics and phages, they do confirm and build upon previous research that observed these synergistic interactions. Our work highlights how using phages can restore the effectiveness of vancomycin against resistant isolates. This finding provides a promising, although unexpected, strategy for moving forward with phage treatments for Vancomycin Resistant Enterococcus infections.

## Introduction

Shortly after the introduction of penicillin and sulfonamides to clinical medicine, antibiotic resistance emerged as a significant issue. Over the past few decades, the prevalence of antibiotic resistant microorganisms has risen at an alarming rate. Unfortunately, currently available antibiotics select for pathogenic bacterial strains with reduced susceptibility[1, 2]. The rapid emergence of antimicrobial resistance (AMR) is a pressing public health issue that poses a significant threat to global health and well-being. The Centers for Disease Control and Prevention (CDC) reported in 2019 that AMR organisms killed at least 1.27 million people globally and approximately 5 million fatalities were associated with AMR pathogens in some manner [2]. The inappropriate use of antimicrobial drugs in humans and animals has been one of the main contributors to the rise of AMR, resulting in a growing number of infections that are difficult and sometimes impossible to treat. AMR is not only a concern for individuals, but also for the healthcare system and economy, as it requires costly and time-consuming measures to develop new drugs and control the spread of resistant bacteria. In the USA alone, antibiotic-resistant bacteria are responsible for 50-60% of hospital-acquired infections [3].

Despite being commensal to the human gastrointestinal and genitourinary tracts, Gram-positive enterococci possess high levels of antimicrobial resistance. As a result, they can cause challenging infections that are associated with mortality rates ranging from 19-48% [4, 5]. These organisms are commonly linked to urinary tract infections arising from medical instrumentation and frequent antimicrobial use. Additionally, they can give rise to infections in the abdominal and pelvic areas, surgical wounds, endocarditis, bacteremia, sepsis, and, albeit rarely, meningitis in newborns [6]. Some enterococci possess intrinsic resistance to commonly employed antibiotics like penicillin and ampicillin. Moreover, they demonstrate elevated resistance to most cephalosporins and all semi-synthetic penicillins due to the presence of low-affinity penicillin-binding proteins [7]. Biofilm formation is an additional important pathogenic feature that may contribute to the creation of bacterial reservoirs that shield pathogens from elimination by antibiotics and increase the risk of morbidity and mortality [8, 9]. *Enterococcus faecalis* and *Enterococcus faecium* are the dominant species of infective enterococci and are responsible for the vast majority of all enterococcal infections in humans [6]. It is noteworthy that *E. faecium* and *E. faecalis* are ranked as the third and fourth most common cause of nosocomial infections globally [3]. The first case of vancomycin-resistant enterococci (VRE) was reported in 1986 from Europe and in 1988 from the USA [10] and later in 1995, the CDC’s Healthcare Infection Control Practices Advisory Committee (HICPAC) classified VRE as a significant emerging pathogen and recommended aggressive infection control measures to prevent its spread [11]. As VRE become more common, the difficulty in treating enterococcal infections has significantly increased. As a result, there is an urgency to develop novel therapeutic approaches [12, 13].

Bacteriophages (“phages” for short) can be used as therapeutic agents to treat multidrug resistant bacterial infections with minimal side effects. Phages are viruses that can infect specific bacterial hosts and replicate within them to produce an abundance of progeny within a short time period before killing their bacterial hosts[14–17]. Phages have not only been shown to be effective in controlling enterococcal growth *in vitro* but also have proven to be successful in treating infections in animal models [18–20]. Likewise, enterococcal phages have been effectively used to disrupt biofilms and treat infections that are usually harder for antibiotics to penetrate [18]. Furthermore, controlled studies showed that unlike antibiotics, phage therapy is highly specific and effective against certain bacterial species without noticeable toxicity or adverse side effects [21–23]. Despite these qualities, the exclusive use of phages to treat bacterial infections suffers from the challenge of treatment-emergent phage resistant bacterial strains [24–27]. Bacteria and phages continuously battle to evolve into a more resilient version of themselves and during this process bacteria acquire/modify their defense mechanisms such as restriction-endonuclease systems, cell surface receptor alteration, CRISPR-Cas9 immunity and abortive infections [28, 29]; thus, bacterial resistance to phages remains a significant consideration.

Phage cocktails can be used as an effective strategy to prevent the emergence of bacteria resistant to phages. In fact, phage cocktails in *in vitro* settings have been shown to be highly effective in controlling the growth of antibiotic and phage-resistant bacterial strains as compared to the use of single phages [30–32]. We have previously shown that a cocktails of two or three phages were effective in limiting the *in vitro* growth of Vancomycin Resistant *Enterococcus faecium* and *faecalis* strains that were originally resistant against some individual phages used in the cocktail [33]. Furthermore, several clinical trial studies have documented successful uses of phage cocktails on a variety of bacterial pathogens [34–36]. Apart from the investigations involving phage cocktails, certain studies have documented encouraging outcomes when exploring the effects of combining phages with antibiotics [31, 37]. A handful of studies in recent years have demonstrated the synergistic effects of phage-antibiotic combinations on vancomycin resistant *E. faecium* [38] and *E. faecalis* [39, 40] strains under various conditions. In the studies reported here, we aimed to confirm and extend prior work to include a greater emphasis on clinical VRE isolates.

The precise molecular mechanisms underlying the enhanced killing of host cells through phage-antibiotic synergistic interactions remain unclear. To address this knowledge gap, we investigated how the combination of phages and antibiotics leads to resensitization of bacterial strains to the antibiotic. Our approach involved comprehensive screening and *in vitro* testing of various phage and antibiotic combinations to identify the most effective synergistic pairs against different enterococcal strains. Additionally, we examined the transcriptome of the bacterial host cells from the most successful phage-antibiotic conditions to identify the key cellular pathways that contribute to these synergistic effects.

## Results

### Characterization of bacteria and phage isolates

Eight isolates of *E. faecium* (Tx1330, EF98PII, EF208PII, and NYU) and *E. faecalis* (DP11, EF116PII, EF140PII, and V587) were identified from patients with infections at UCSD Health or were type strains (**Table 1**). These isolates were chosen on the basis of their vancomycin (VAN) susceptibility profiles. Isolates Tx1330 and DP11 were vancomycin susceptible, while EF98PII, EF208PII, NYU, EF116PII, EF140PII, and V587 were vancomycin resistant (**Table 1**). Some of these isolates were sequenced as part of this study, however, some had previously undergone whole genome sequencing [33].

**Table 1.** Enterococcus isolates used in this study and their antibiotic susceptibility profiles, and the phages used for synergy experiments. The EUCAST System for Antimicrobial Abbreviations was used to name antibiotics. GEN: Gentamycin, VAN: Vancomycin, TET: Tetracycline, CZO: Cefazolin, CXI: Cefoxitin, CLI: Clindamycin, ERY: Erythromycin, GEN: Gentamicin, GEN-Syn, LEV: Levofloxacin, MIN: Minocycline, STR-Syn: Streptomycin synergy, TET: Tetracycline, TRS: Trimethoprim-Sulfamethoxazole.

| Enterococcus strains |  | GenBank Accession | Vancomycin sensitivity | Antibiotic Resistance | Phages used for PAS study |
| --- | --- | --- | --- | --- | --- |
| *E. faecium* | Tx1330 | GCA_003583905.1 | S | CZO, CXI, GEN, CLI, TRS | Bop |
|  | EF98PII | SAMN39584152 | R | VAN, AMP, BEN, TET | Ben |
|  | EF208PII | SAMN36748517 | R | VAN, CZO, CXI, CLI, ERY, GEN, LEV, MIN, Penicillin G, STR-Syn, TET, TRS | Bop |
|  | NYU | SAMN36748518 | R | VAN, AMP, CZO, CXI, GEN, CLI, TRS, ERY, LEV, MIN, MOX, BEN, TET | Ben |
| *E. faecalis* | DP11 | JALPNV000000000 | S | GEN | ReUmp |
|  | EF116PII | SAMN39584151 | R | VAN, GEN, TET | PL |
|  | EF140PII | SRX21185070 | R | VAN, CZO, CXI, CLI, ERY, GEN, GEN-Syn, LEV, MIN, STR-Syn, TET, TRS | Bob |
|  | V587 | GCA_000394175.1 | R | VAN, GEN, ERY, CZO, CXI, CLI, TRS | Ben |

We also characterized phages active against many of the *E. faecium* and *E. faecalis* isolates we found. Some of these phages had previously been characterized in our prior studies [33]. We obtained the phage genome sizes using whole genome sequencing (**Table 2**), and morphologies were obtained using transmission electron microscopy (TEM; **Figure 1, panels A-E**). The TEM confirmed phage morphologies consistent with myoviruses and siphoviruses, respectively (**Figure 1**). Specifically, phages Ben, Bob, and Bop displayed large icosahedral heads with medium sized contractile tails consistent with myovirus morphologies (**Figure 1, panels A-C**), while PL and ReUmp exhibited prolate shaped heads with noncontractile tails, consistent with siphovirus morphologies (**Figure 1, panels D-E**).

**Figure 1.**
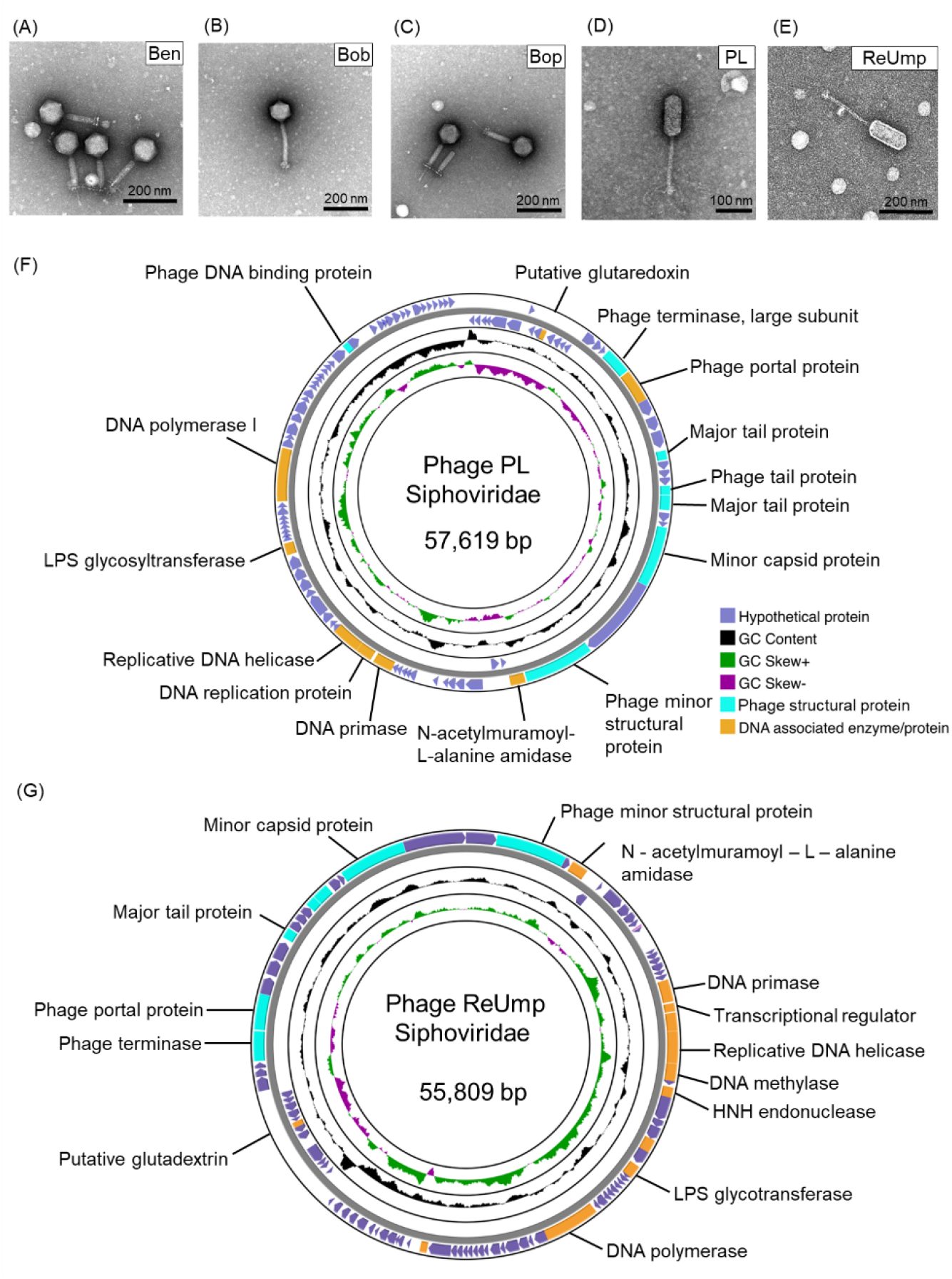
Morphological and genomic characterization of *E. faecium and E. faecalis* phages. Myovirus morphologies were observed for phages Ben, Bob and Bop (A-C). Siphovirus morphologies were observed for phages PL and ReUmp (D-E). Genome maps of phage PL (F) and ReUmp (G). The outermost circle shows open reading frames (ORF) of predicted proteins. Phage structural proteins are highlighted in cyan; DNA associated proteins are shown in yellow, hypothetical proteins are shown in purple.

**Table 2.** Enterococcus phages used in this study. Genome sizes were obtained from whole genome sequencing. Morphologies were obtained via TEM.

| Phage | Genome size (bp) | Morphology | Isolation source | Reference |
| --- | --- | --- | --- | --- |
| Ben | 151,985 | Myoviridae | Orange county sewage, CA | Wandro et. al., 2022 |
| Bob | 142,921 | Myoviridae | Orange county sewage, CA | Wandro et. al., 2022 |
| Bop | 153,454 | Myoviridae | Orange county sewage, CA | Wandro et. al., 2022 |
| PL | 57,619 | Siphoviridae | Point Loma sewage, CA | This Study |
| ReUmp | 55,809 | Siphoviridae | Orange county sewage, CA | This Study |

### A liquid assay to demonstrate phage-antibiotic synergy (PAS)

We developed a comprehensive screening assay to determine optimum phage and antibiotic concentrations that could efficiently inhibit the growth of Vancomycin Resistant Enterococcus (VRE) and Vancomycin Susceptible Enterococcus (VSE) isolates. To perform this assay, we set up 96-well plates with increasing concentrations of antibiotics across the x-axis and increasing concentrations of the phages across the y-axis (**Figure 2**). We examined three different antibiotics which are commonly used against enterococcus isolates, including vancomycin (VAN), linezolid (LZD), and ampicillin (AMP). Of note, many of the enterococcus isolates (both *E. faecium* and *E. faecalis*) were resistant to vancomycin, while the *E. faecium* isolates were considered intrinsically resistant to ampicillin (**Table 1**). The three antibiotics were used in combination with the 5 different phages that exhibited lytic activity against the enterococcus isolates included in this study.

**Figure 2.**
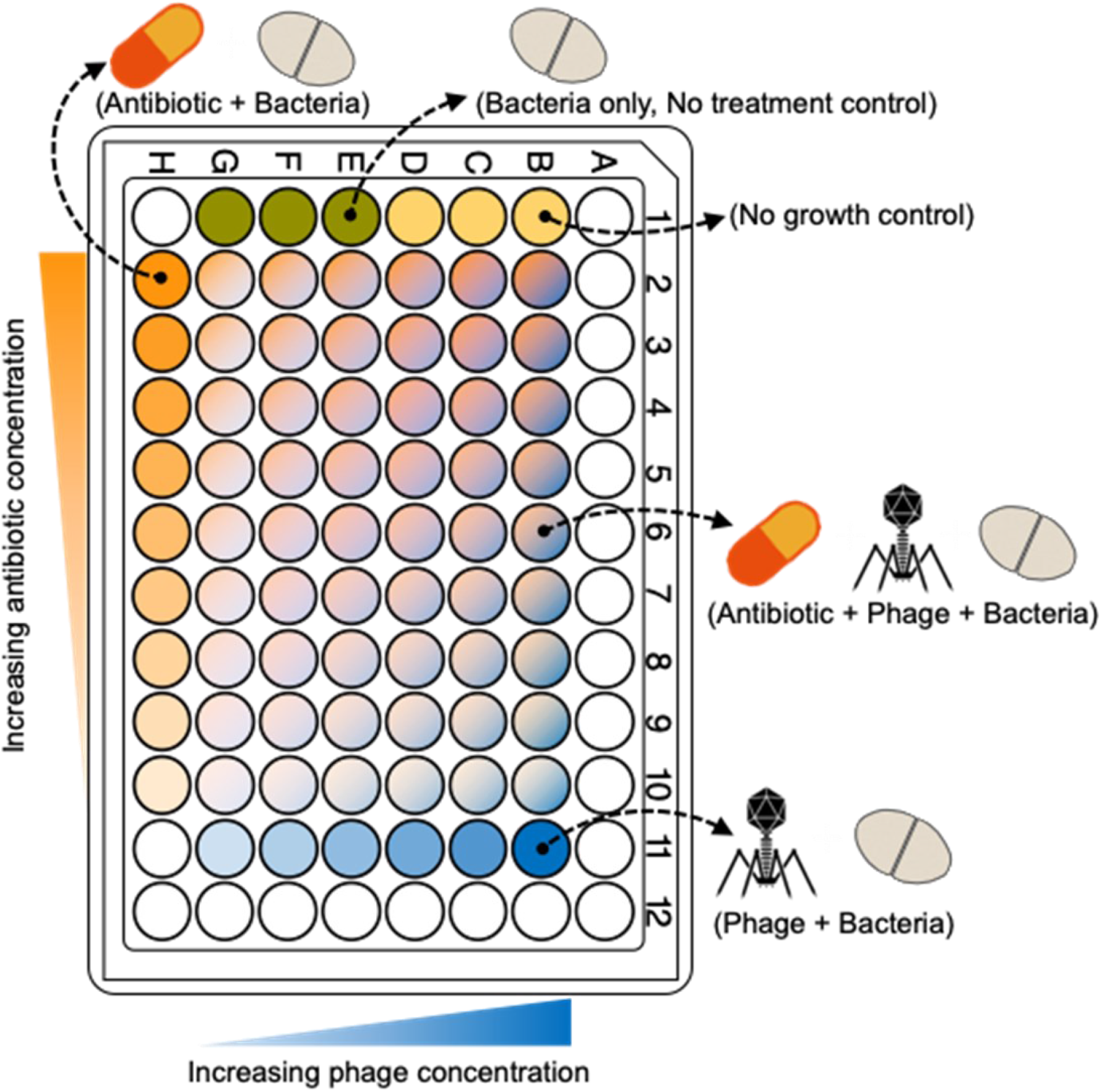
Experimental layout of the phage-antibiotic co-cultivation assay. Bacterial growth in the presence of antibiotics, phage, or both were studies using a 96 well plate. Differently colored wells represent various treatments: Antibiotic, phage, antibiotic + phage, no treatment control, and no growth controls. Color intensity in the individual wells represent their respective concentration gradient. Some outer wells (colorless) bordering the plate were unused and were filled with BHI media.

In each assay, a logarithmically growing enterococcal culture was incubated with a series of different concentrations of phages and antibiotics. Bacterial growth was monitored for 18 hours by measuring the OD_600_ (**Figure 2**). The results were represented by Phage-antibiotic synergy (PAS) diagrams, which have previously been termed synograms by Liu et al. [46], where the percentage bacterial growth reduction is represented by a color gradient. We developed PAS diagrams for *E. faecium* clinical isolates EF98PII and NYU for each of the aforementioned antibiotics and phage Ben. As previously mentioned, both *E. faecium* isolates are intrinsically resistant to ampicillin, but also express high level resistance to vancomycin (VRE). When co-cultivated with phage Ben, both isolates demonstrated a growth reduction between 80-90% at relatively minimal levels of the antibiotics (1-2 µg/ml of vancomycin and 0.25-0.5 µg/ml of ampicillin) (**Figure 3B and 3D**). Similar results were observed for linezolid with phage Ben, where significant reductions were observed in the antibiotic MIC’s when phage Ben was added (**Figure 3B and 3D**). We observed a moderate growth reduction when laboratory adapted *E. faecium* isolate Tx1330 was used with phage Bop for each of the antibiotics used (**Figure 3A**). Minimal growth reduction was observed for the VRE clinical isolate EF208PII, phage Bop and each of the antibiotics tested (**Figure 3C**). The optimal phage titer that exhibited PAS in co-cultivation experiments was in the range of 10^4^-10^6^ phages/mL for all the experiments. The growth reduction was significantly lower (one way ANOVA, p<0.001) in the PAS wells than in wells with only antibiotics or phages (**Figure S2 and Table S1**). Our data show significant growth reductions for both VRE and VSE isolates of *E. faecium* when phages are combined with antibiotics regardless of whether the enterococcus isolate has prior resistance to the antibiotic in question. High resistance levels to vancomycin did not affect whether we observed synergistic interactions between phages and vancomycin for these isolates.

**Figure 3.**
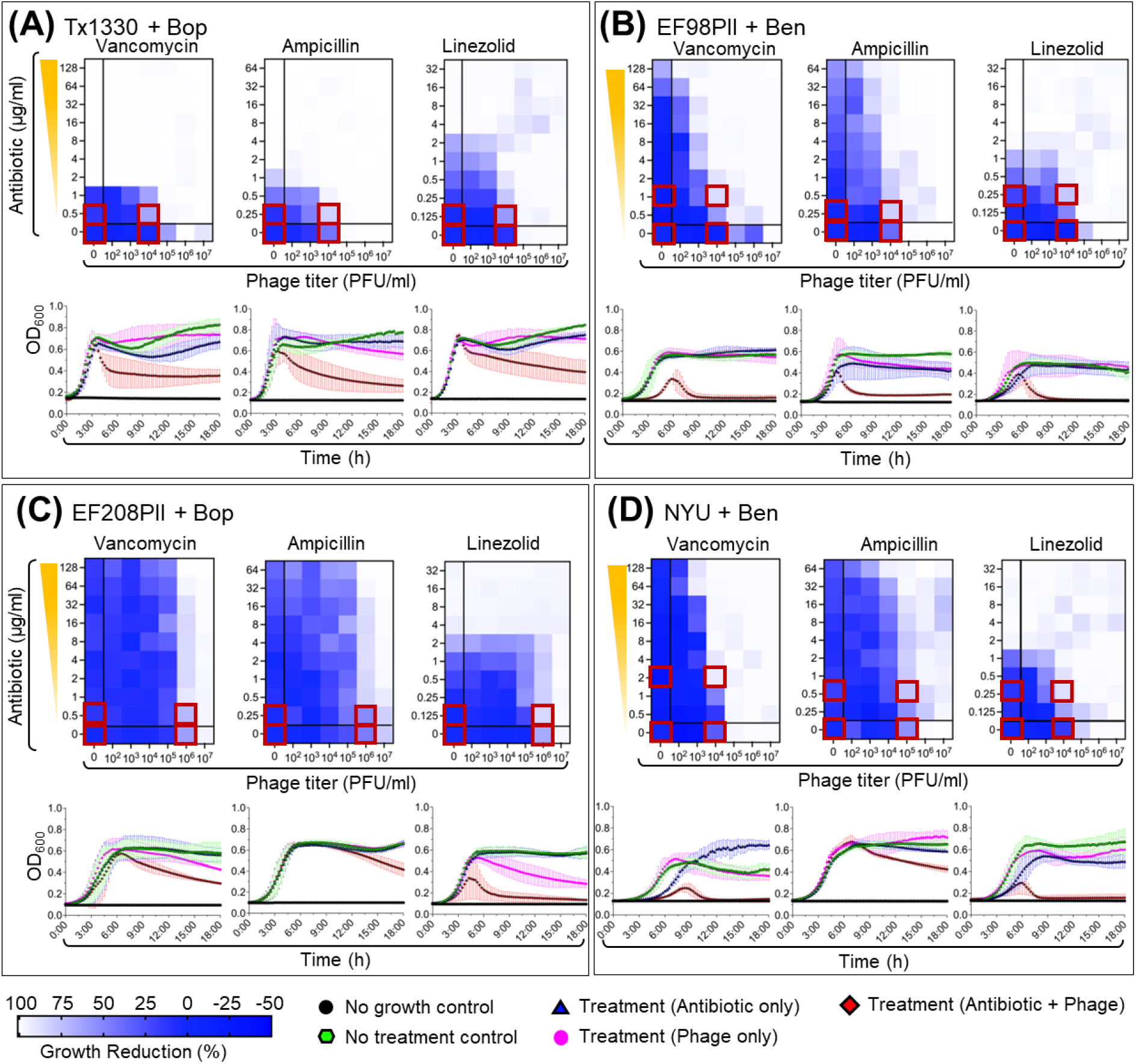
Synograms and growth curves showing the effect of various treatments (antibiotics, phages, antibiotic + phages) on growth dynamics of *Enterococcus faecium*. The color gradient in synograms represents the percentage of growth reduction. Growth reduction was calculated using following formula: Percentage reduction = [(OD growth control –OD treatment)/OD growth control] * 100. The average of three biological replicates is shown. Antibiotics vancomycin, ampicillin and linezolid were tested in all strains. A) *E. faecium* Tx13301 in the presence of phage Bop. B) *E. faecium* EF98PII in the presence of phage Ben. C) *E. faecium* EF208PII in the presence of phage Bop. D) *E. faecium* NYU in the presence of phage Ben

While *E. faecium* is the most commonly observed VRE isolate in clinical medicine [41], *E. faecalis* often is capable of acquiring the same mobile genetic elements and becoming VRE. Because of the significant genetic differences between *E. faecium* and *E. faecalis* isolates, we next tested whether we observed similar trends in PAS for *E. faecalis* that we observed for *E. faecium*. We examined two separate clinical isolates EF116PII and EF140PII, along with two laboratory-adapted isolates DP11 and V587. Isolates EF116PII, EF140PII, and V587 were resistant to each of the antibiotics tested (vancomycin, ampicillin, and linezolid), while DP11 was susceptible to them all. We first tested VRE isolate EF116PII with phage PL and found that it was almost completely inhibited when only very low concentrations of vancomycin (1 µg/mL) were added (**Figure S1, Panel B**). There also were moderate improvements in the inhibition of this isolate in the presence of relatively low concentrations of ampicillin and linezolid. Many of these results were statistically significant (**Figure S2, Panel F**). We also identified significant results for *E. faecalis* isolate DP11 in the presence of vancomycin and phage ReUmp (**Figure S1, Panel A**), EF140PII in the presence of ampicillin and phage Bob (**Figure S1, Panel C**), and V587 in the presence of all antibiotics tested and phage Ben (**Figure S1, Panel D**). Many of these results were statistically significant, indicating that the synergistic interactions demonstrated between antibiotics and phages were robust (**Figure S2, Panels E-H**). These results indicate that many of the results observed with *E. faecium* were reproducible for *E. faecalis* and suggest a trend with enterococcus where synergistic interactions may be observed between antibiotics and phages regardless of whether pre-existing susceptibility to antibiotics is present.

### Transcriptomes from *E. faecium* and *E. faecalis* in PAS conditions

We sought to characterize the transcriptomes of the *E. faecium* and *E. faecalis* isolates to discern whether there were differences in gene expression profiles that might account for the synergistic responses to the phages and antibiotics. We compared the gene expression profiles in response to phages and antibiotics with those of antibiotics alone, phages alone, and with growth controls. Each of the enterococcus isolates was co-cultivated with specific phages, one of three different antibiotics, including vancomycin, linezolid, and ampicillin, or no treatment.

Because we needed relatively large quantities of RNA for transcriptome sequencing, we reproduced the experiments for *E. faecium* and *E. faecalis* at larger volumes on the conditions that were already identified as producing synergy. Three biological replicates for each of the enterococcus isolates and for all phage and antibiotic combinations were obtained. Transcriptomes were obtained at 5 and 18 hours post treatment (**Figure S3-A**), and the number of reads ranged from 2.5 × 10^7^ to 1.0 × 10^8^. There were generally fewer reads recovered at 18 hours compared to 5 hours, which may correspond to fewer recovered cells at the later time point (**Figure S3-B**).

We next clustered specimens together based on their transcriptome profiles to decipher whether patterns emerged based on antibiotics used, phages used, or the combination of both. Specimens were clustered together using Principal Component Analysis (PCA) and were labeled according to time, antibiotic, phage treatment, and no treatment (**Figure S4**). There were clear differences in expression profiles in both *E. faecalis* and *E. faecium* isolates when comparing gene expression at 5 hours and 18 hours, but there were not obvious segregation of antibiotic and antibiotic/phage groups as shown by overlapping ellipses.

Next, we segregated the isolates based on sampling time and performed PCA analysis by treatment groups to discern whether we could identify differences in gene expression profiles in phage, antibiotic, and dual antibiotic/phage synergy treatment groups (**Figure 4**). For *E. faecium* EF98PII, there was significant variation among the different treatment groups at both 5 and 18 hours with PC1 of 53% and 59%, respectively; similar results were identified for *E. faecalis* EF116PII. There was distinct clustering identified when comparing the antibiotic groups to the antibiotic/phage groups at 18 hours, but not at 5 hours for both the *E. faecium* and *E. faecalis* isolates (**Figure 4**). This was true regardless of the antibiotic used, but the largest segregation was observed for vancomycin and linezolid, with generally less segregation observed for ampicillin. Phage only and growth control groups were found to be colocalized on the plot irrespective of sampling time.

**Figure 4.**
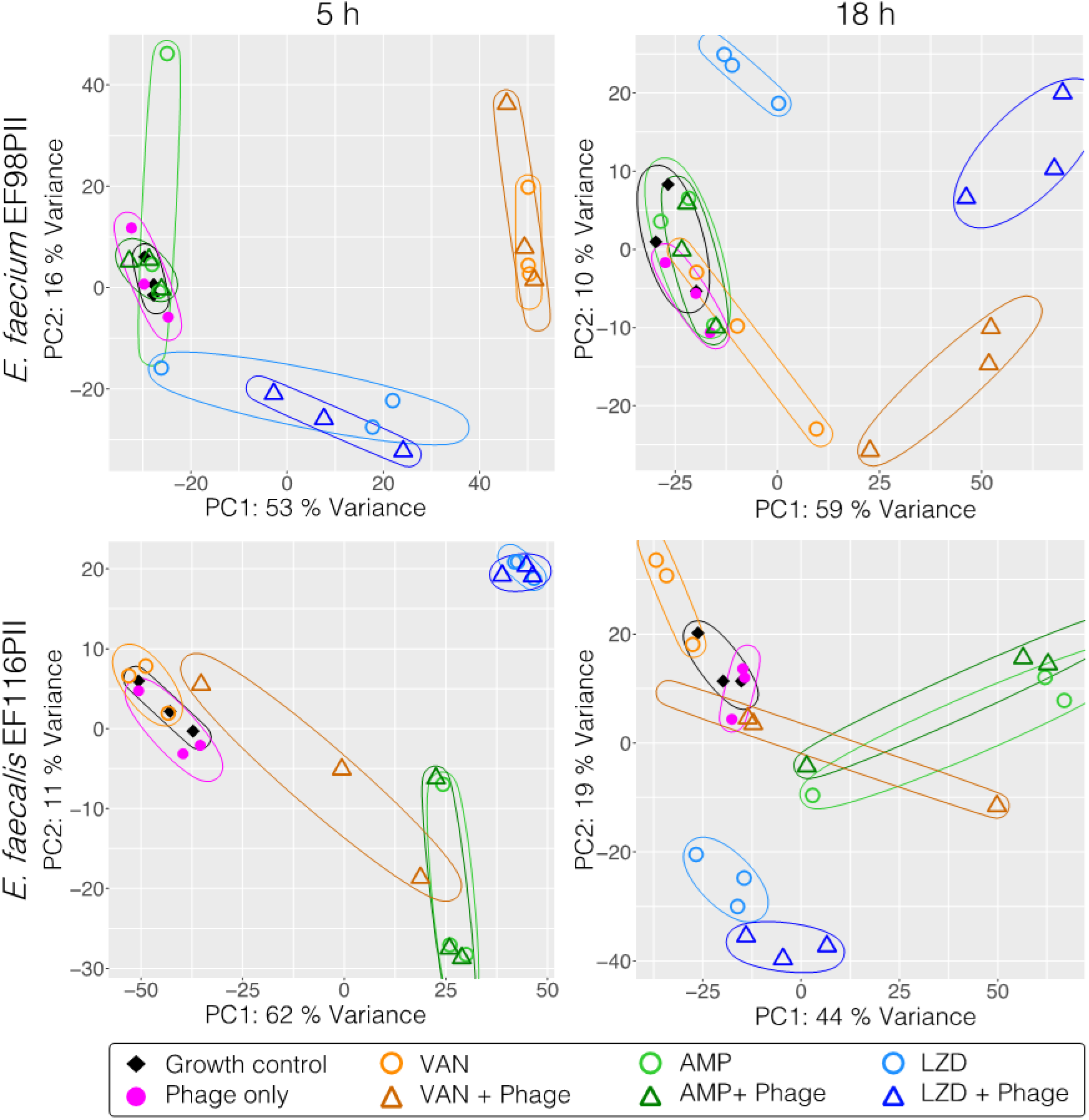
Principal component analysis (PCA) showing clustering of samples at different time points and pathway analysis of PCA rotation. (A) Scatter plots of the first two principal components of the normalized gene expression profile of all the samples. The ellipse encircles the three biological replicates for individual experimental conditions and were drawn at tolerance cutoff of 0.01.

### Phage-antibiotic synergy alters cellular stress responses and membrane transport

Because there was clear delineation in gene expression profiles when comparing antibiotics versus the combined effects of antibiotics and phages at 18 hours compared to 5 hours, we focused our analysis on the 18-hour time point moving forward to identify differences that might account for the synergistic effects observed. We performed differential gene expression analysis using iDEP v0.96 to identify the top 50 differentially expressed genes between the antibiotic only and the antibiotic/phage synergy. We found that the majority of top differentially expressed genes were involved in membrane transport, DNA replication and damage repair, transcription regulation, and cellular stress response regulation (**Figure 5 and Table S2 and Table S3**). There is a distinct separation between the highly expressed and minimally expressed genes in the vancomycin and linezolid treatment groups, while the distinction is less in the ampicillin treatment group (**Figure 5**).

**Figure 5.**
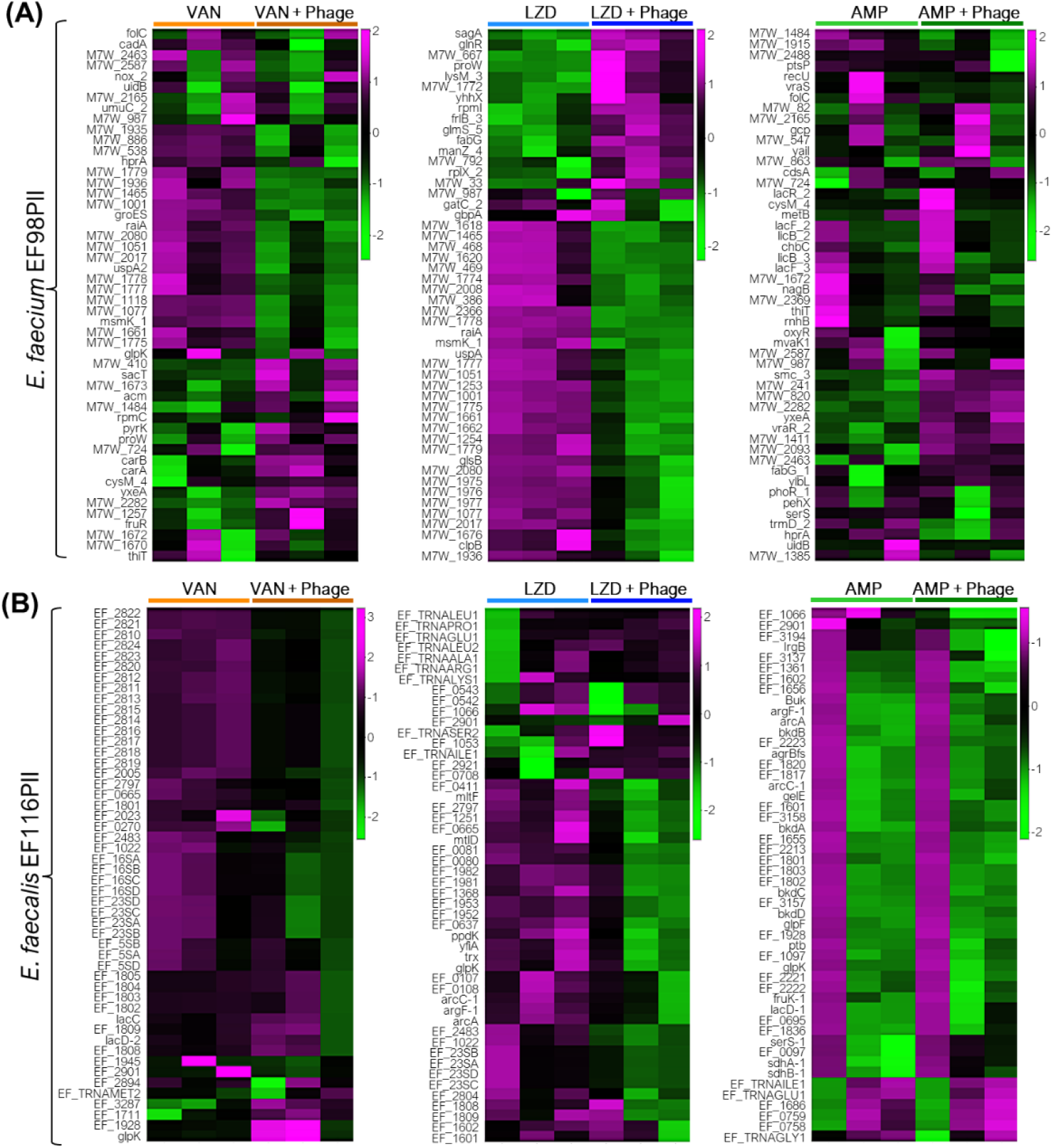
Heatmap showing top 50 differentially expressed genes. in *E. faecium* EF98PII (A) and *E. faecalis* EF116PII (B) for various comparison between “Antibiotic + Phage” versus “Antibiotic” only treatment samples at 18 h post co-culture. The data were centered by subtracting the average expression level for each gene and samples were normalized by dividing with SD. Pearson’s correlation coefficient and average linkage was used to calculate distance matrix. Each sample group includes three biological replicates and is highlighted by colored bars atop the heatmaps. The color key legend next to each heatmap represents the degree of variation in the expression of genes between samples.

In the presence of vancomycin and phages in *E. faecium* isolate EF98PII, some of the highly expressed genes encode proteins involved in membrane transport systems.

Some of those genes include: Cadmium-translocating P-type ATPase (*cadA*), manganese transport protein (mntH), ABC transporter ATP-binding protein (*msmK_1*), substrate binding domain of ABC-type glycine betaine transport system (*proW*), PTS system (fructose-specific IIa, IIB, IIC components), hydroxymethylpyrimidine ABC transporter (transmembrane component) and energy-coupled thiamine transporter ThiT (*thiT*). Similarly, heat shock protein Hsp20, universal stress protein UspA2, and co-chaperone (*groES*) involved in regulating stress responses and protein folding were among the top 50 differentially expressed genes (**Table S2**). Additionally, genes involved in membrane transport systems, stress response and ribonucleoprotein and protein biosynthesis were found to be expressed under both ampicillin/phage and linezolid/phage treatment conditions in *E. faecium* EP98PII (**Table S2**).

For *E. faecalis* isolate EF116PII, we identified many of the identical sets of highly differentially expressed genes that were also identified in *E. faecium* isolate EF98PII (**Tables S2 and S3**), indicating that despite the significant genetic differences between the two species, the responses to the antibiotics and phages were highly similar. We also identified additional differentially expressed genes in *E. faecalis* EF116PII. Some of these were related to prophages, as they represented a major capsid protein, minor structural protein, tail tape measure protein, and major tail protein, and were all downregulated in response to the combination of vancomycin/phage (**Figure 5B and Table S3**).

We next used DeSeq2 to characterize differentially expressed genes between the antibiotic and the phage/antibiotic conditions to determine whether similar results were identified by different methods and whether additional differentially expressed genes could be identified. We used a minimum fold change value of 2 with a minimum FDR of 0.05. We found that a total of 352/329 and 304/210 genes were significantly (Padj < 0.05) upregulated/downregulated in *E. faecium* EF98PII and *E. faecalis* EF116PII, respectively, when the vancomycin/phage treatment group was compared to the vancomycin group. Similarly, 464/465 and 10/34 genes were found to be significantly (Padj < 0.05) upregulated/downregulated in *E. faecium* EF98PII and *E. faecalis* EF116PII respectively in the linezolid/phage treatment compared to the linezolid group (**Figure 5A**). However, as suggested by the low degree of variability between ampicillin/phage and ampicillin groups (**Figure 4**), we see a relatively small pool of genes significantly (Padj < 0.05) upregulated/downregulated (11/1) in *E. faecium* EF98PII and none in the case of *E. faecalis* EF116PII (**Figure 5A**).

There were a number of different genes identified in *E. faecium* EF98PII using DeSeq2 that were highly upregulated under vancomycin/phage synergistic conditions compared to vancomycin alone (**Table S4**). These include a transcriptional repressor of the fructose operon (M7W_1255, fruR), a transcription anti-terminator (M7W_2638, *sacT*), an oligopeptide transport system permease protein (M7W_2290; *oppB*), a PTS system, fructose-specific component (M7W_1257), and sortase A (M7W_698) among others (**Table S4**). There also were a number of significantly downregulated genes in vancomycin/phage synergistic conditions compared to vancomycin alone. These included a number of hypothetical genes, organic hydroperoxide resistance (M7W_1936), heat shock protein Hsp20 (M7W_2017), multiple sugar ABC transporter, manganese transport protein MntH (M7W_1001), universal stress protein family (M7W_1568, *uspA2*), ATP-binding protein (M7W_2275, *msmK_1*), cadmium-translocating P-type ATPase (M7W_1465), and others (**Table S4**)

We also examined those genes that were differentially regulated in *E. faecium* EF98PII at 18 hours using DeSeq2, focusing on those that were significantly upregulated under linezolid/phage synergistic conditions compared to linezolid alone. These genes included a D-serine, D-alanine, glycine transporter (M7W_814), sortase A (M7W_73), oligopeptide transport system permease (M7W_2290, *oppB*), ABC transporter membrane-spanning permease glutamine transport (M7W_2302, *yecS*), and a glycine betaine ABC transport permease (M7W_2392, *proW*) among others (**Table S4**). We also identified some downregulated genes, which included heat shock protein Hsp20 (M7W_2017), an N-acetylglucosamine-specific IIA, IIB, IIC component (M7W_1488), a cadmium-translocating P-type ATPase (M7W_1465), a manganese transport protein MntH (M7W_1001), a universal stress protein family (M7W_1773, *uspA*), a multiple sugar ABC transporter, an ATP-binding protein (M7W_2275, *msmK_1*), an abortive infection protein (M7W_2008), a universal stress protein family (M7W_469), and a putative hydrolase of alpha, beta superfamily (M7W_1662), among others (**Table S4**).

When examining the effects on *E. faecium* EF98PII of ampicillin/phage synergy compared to ampicillin alone at 18 hours, we identified fewer genes that had altered regulation compared to linezolid and vancomycin (**Table S4**). We identified upregulation in a transporter associated gene associated with vraSR (M7W_1411) and a response regulator (M7W_1413, *vraR_2*). We identified downregulation in other hypothetical genes (**Table S4**).

We also characterized *E. faecalis* EF116PII using DeSeq2 to identify overlapping upregulated or downregulated genes between *E. faecalis* (EF116PI) and *E. faecium* (EF98PII) during phage and antibiotic selection. We began by examining those genes that were significantly upregulated under vancomycin/phage synergistic conditions compared to vancomycin alone (**Table S5**). Those genes included an ABC transporter ATP-binding protein (EF_1333), a site-specific integrase (EF_0479), and an amino acid permease (EF_1103). Downregulated genes included a phage associated holin (EF_2803/2804), a phage tail protein (EF_2005), a phage baseplate upper protein (EF_2810), a HK97 gp10 family phage protein (EF_2007), a PTS transporter subunit EIIC (EF_0270), and a PTS sugar transporter subunit IIA (EF_0412, *mltF*), among others.

We next examined the gene expression responses of *E. faecalis* EF116PII to linezolid/phage synergy compared to linezolid alone (**Table S5**) using DeSeq2 to identify upregulated and downregulated genes. We identified many upregulated genes in response to linezolid/phage synergy that mostly represented transcriptional regulators and ABC transporters. These genes included ABC transporter ATP-binding protein (EF_2652, *potA*), ABC transporter substrate-binding protein (EF_2649), ABC transporter permease (EF_2650/2651), ABC transporter ATP-binding protein (EF_1673), Sigma-54-dependent transcriptional regulator (EF_1010), and response regulator transcription factor (EF_0926), among others (**Table S5**). Other genes were downregulated in response to linezolid/phage synergy, which included mostly membrane transporters, phage associated genes, toxin-antitoxin systems, and sugar transporters. These genes included PTS sugar transporter subunit IIA (EF_0412, *mltF*); PTS mannitol transporter subunit IICBA (EF_0411), phage associated protein holin (EF_2803), phage tail protein (EF_2001/2005), phage baseplate upper protein (EF_2810), phage tape measure protein (EF_2003), type II toxin-antitoxin system RelB/DinJ family (EF_0512), and type II toxin-antitoxin system YafQ family (EF_0513).

### Pathway analysis

We next used the STRING database [42] to identify biosynthetic pathways that may be involved in antibiotic/phage synergy responses. We first evaluated the response in *E. faecium* EF98PII and found that pathways involved in purine and pyrimidine biosynthesis were significantly enriched in both the vancomycin/phage (**Figure 6-A)** and linezolid/phage (**Figure 6-B**) responses compared to antibiotics alone. In contrast, we identified various stress response pathways that were inhibited, along with suppression of chaperones, membrane transport channels, and iron-sulfur cluster biosynthesis. In response to ampicillin/phage synergy, fewer pathways were identified. These included a small number of upregulated genes involved in cell-wall biosynthesis, phage shock protein C, and transcription regulators such as Helix-turn-helix domain *rpiR*.

**Figure 6.**
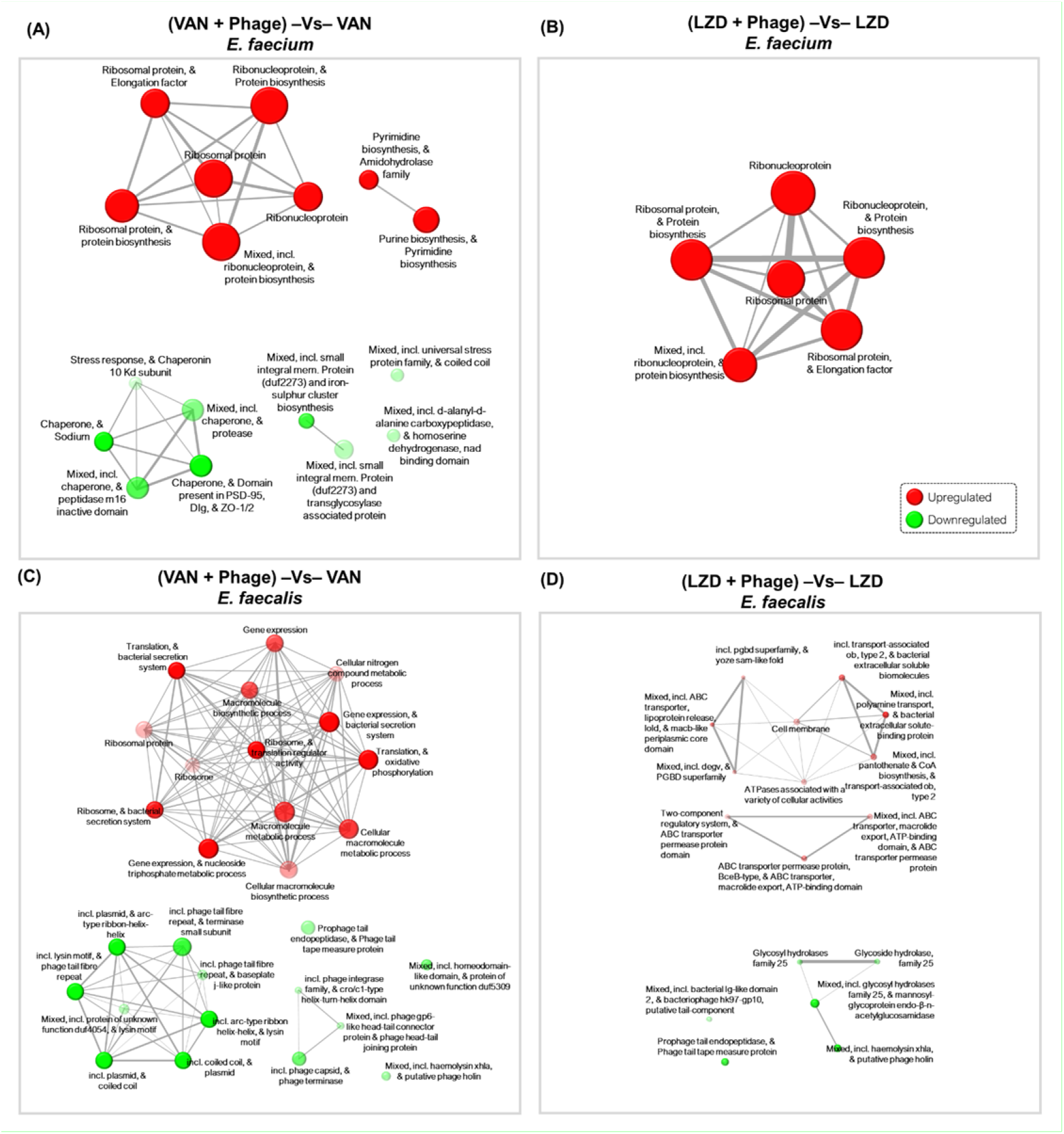
Pathway analysis of significantly upregulated/downregulated genes during “Phage +Antibiotic” co-culture as compared to “Antibiotic” only culture at 18 h. Network of pathways involving differentially expressed genes isolated from *E. faecium* EF98PII (A,B) and *E. faecalis* EF116PII (C,D) treated with (Antibiotic + Phage) versus “Antibiotics” only at 18 h. Genes were filtered as background and “all available gene sets” were used for enrichment analysis of differentially expressed genes. Two pathways (nodes) are connected if they share 30% or more genes. Red represents upregulated, green represents downregulated. The node sizes correspond to their respective gene set sizes. Darker colored nodes correlate to more significantly enriched gene sets. Thicker edges represent more overlapped genes.

We also used the STRING database to identify biosynthetic pathways that may be involved in the responses to antibiotic/phage synergy in *E. faecalis* EF116PII. Most of the gene pathways being downregulated when vancomycin/phage was present were associated with phages, while biosynthetic pathways were significantly enhanced (**Figure 6-C**). In response to linezolid/phage synergy PGBD-like superfamily gene pathways involved in peptidoglycan binding, and ABC transporter permease pathways that confer antibiotic resistance in gram-positive bacteria were upregulated (**Figure 6-D**).

## Discussion

There is a significant need to identify alternatives for the treatment of enterococcal infections in humans. The organism has the potential to resist many commonly used antibiotics through the acquisition of a plasmid (such as vanA or vanB), a transposon, or through the presence of genes on the genome (such as vanD) in certain species. Although vancomycin has historically been a critical component of most enterococcal treatment regimens, an increasing prevalence of VRE over the past decade has mandated the development of alternative treatment approaches. Patients who have received extensive antibiotic therapy including bone marrow and organ transplant recipients, are chronically ill, or are in long-term hospital care facilities, are at particular risk from VRE infections. In these patients, the enterococci can be extremely difficult to eradicate and often recur following cessation of therapy. Thus, alternative treatments, such as phage therapy, and protocols for decolonization of these patients of their enterococcus infections have become increasingly attractive options.

Phages have become alternative and adjunctive therapies to antibiotics for multi-drug resistant bacteria but have been difficult to deliver in a timely fashion to patients in acute clinical situations. Part of this has been due to the manner in which phages are evaluated to determine whether they may be efficacious against the pathogens that are associated with disease severity. Generally, pathogens such as enterococci are identified causing an illness, and then those pathogens are referred to centers that are willing to screen for phages that are capable of lysing those pathogens. Generally, those centers will identify whether those phages are capable of killing the pathogens with efficiency of plating values [43] This procedure generally ignores the fact that the phage and or phages that will be delivered to the patient will almost never be delivered alone without the concomitant delivery of standard-of-care antibiotics. Thus, we sought to characterize a more realistic situation that more accurately reflects the situation that is observed clinically when patients receive phage therapy. That is, what is observed when the patient receives both phages and antibiotics simultaneously.

We evaluated the impact of phages and antibiotics when used concurrently extended and confirmed prior observations. Those findings were that antibiotics such as vancomycin could work synergistically with phages to eliminate the enterococci [41, 44] Our results extended those findings by demonstrating that this was true for both *E. faecium* and *E. faecalis*, for clinical isolates and for laboratory adapted isolates of enterococci, and most importantly, that synergy was observed whether or not the enterococcal isolate was antibiotic susceptible. For example, we observed significant synergistic interactions with ampicillin and phages in *E. faecium*, which is a combination that would almost never be used clinically because *E. faecium* is considered to be intrinsically resistant to ampicillin [45]. Perhaps more importantly, we confirmed in our clinical isolates of VRE *E. faecium* and *E. faecalis* that despite their confirmed high-level resistance to vancomycin (**Table 1**), when combined with phages, the MICs to vancomycin were reduced to within readily achievable ranges (**Figure 3**). Such a finding has significant implications for the clinical treatment of recalcitrant VRE infections as well as decolonization protocols for VRE, as it indicates that antibiotics that had previously been deemed useless against this MDR pathogen may have significant utility when used in combination with phages.

It is important to note that the antibiotic resistance in this study that was overcome through the process of synergy was not a permanent phenomenon. Indeed, we tested each of these isolates afterwards just to demonstrate that they were still antibiotic resistant (data not shown) despite the fact that they had susceptible MICs when they were tested in the presence of both phages and antibiotics. This was important because it indicated that the combined effect of the antibiotic and the phage required the ongoing presence of phages. Upon removal of the phages, the high-level ampicillin or vancomycin resistance returned. This is contrary to a situation that has been observed with other phages with organisms such as *Pseudomonas aeruginosa* [38], where the utilization of certain phages leads to more permanent changes in antimicrobial susceptibilities that are not reversed with the removal of the phages.

We expected that we might find combined effects of phages and antibiotics when we treat enterococci with them both together. Indeed, we identified these phenomena when we developed a solid media assay for assessing the combined effects of antibiotics and phages on multiple different bacteria, including *E. faecium* and *E. faecalis* [46] What is surprising to us is the ability to overcome these phenomena in clinical VRE isolates that are confirmed to have *vanA* expressing plasmids. These plasmids are known to express high levels of the vanA genes that result in the proteins saturating the cell walls of the enterococcal cell surfaces, resulting in high levels of resistance to vancomycin, which no longer bind to these altered VanA [47]. We did not expect to essentially restore vancomycin susceptibility in these organisms through the addition of a phage, as one might assume the restoration might occur through the downregulation of vanA (which was not observed), or the upregulation of the native gene (which also was not observed). Thus, a complex combination of regulatory factors involving nucleotide and protein biosynthesis, stress responses, and transport appear to be involved in the synergy responses rather than just a reversal of the original antibiotic resistance response. While we could not pinpoint the exact mechanism responsible for the restored susceptibility to the antibiotic and phage using transcriptomic analysis, a study such as this one is just the first step in elucidating what may be necessary to pinpoint mechanism behind developing synergistic responses between antibiotics and phages.

## Materials and Methods

### Bacterial strains, growth conditions and quantification

Enterococcus isolates used in this study were collected from patients admitted at the University of California San Diego (UCSD) Health. Four different strains each of *E. faecium* and *E. faecalis* were used to study the synergistic interaction between antibiotics-phages and their antibiotic resistance profiles were determined using broth microdilution techniques using BD Phoenix instrument (Becton Dickinson, Franklin Lakes, NJ, USA) at the UCSD Centre for Advanced Laboratory Medicine (**Table 1**). For each species we tested three VAN resistant and one sensitive strain against a combination of their respective phages with antibiotics: (Vancomycin (VAN), Ampicillin (AMP), and Linezolid (LZD)) or antibiotics only to screen for antibiotic-phage synergy treatment condition (Table 1). All bacterial strains were cultured in Brain Heart Infusion (BHI) medium (BD Difco™, Catalog# DF0418-17-7) at 37°C with shaking at 200 rpm (for liquid culture) and supplemented with antibiotics during synergy experiments. For bacterial propagation and maintenance, solid BHI media were used with 1.5 % agar while for plaque assays 0.4% top agar were used over the solid media. For quantification, 100 µl of diluted bacterial samples from each treatment condition (usually dilution 10^-4^, 10^-5^ & 10^-6^) was spread onto 1.5% BHI agar plates and incubated overnight at 37 °C. The following day, the number of colonies were counted on each dilution plate and the colony forming units per ml (CFU/ml) for each sample was determined.

### Bacteriophages isolation, propagation, quantification, and characterization

The phages used in this study (**Table 2**) were isolated from sewage samples from different regions of southern California using multiple enrichment protocol as described elsewhere with some modifications [48]. Briefly, sewage samples were centrifuged at 10,000 × g for 10 minutes at 4°C to remove particulate matters and then 20 ml supernatant was mixed with an equal volume of double strength (2×) BHI broth. To this mixture, 500 μl of overnight grown culture of *E. faecium* or *E. faecalis* (maintained at OD_600_ ∼ 0.2) was added and incubated overnight at 37°C in a shaker incubator. One percent v/v chloroform was added to the mixture, vortexed and incubated at room temperature (RT) for 30 minutes. This was followed by centrifugation at 5000 × g for 15 minutes at 4°C and supernatant was filtered through sterile 0.45 μm PVDF syringe filters (Whatman Puradisc, Item# 6746-2504). Next, to screen for phages against these bacterial species, spot assays were performed as follows. One hundred μl of overnight grown culture of enterococcus (diluted to OD_600_ ∼ 0.2) were mixed with molten BHI top agar (cooled to ∼ 45°C) and uniformly spread over the BHI agar plates. After the soft agar was solidified, 5 μl of filtered phage lysate (from different sewage samples) was spotted onto their respective bacterial species plates. Spots were air dried followed by overnight incubation of plates at 37°C. Plates were examined for clear spots and if positive, were further processed for three rounds of phage purification by plaque assay as described by Wandro *et. al.* [33]. Briefly, clear spots were picked by 1 ml sterile pipette tip and resuspended in 100 μl sterile PBS buffer. Following brief vortexing and centrifugation, the supernatant was mixed with equal volume of *Enterococcus sp.* culture (OD_600_ ∼0.2) and incubated at 37°C for 10 minutes. Molten BHI top agar (5 mL; cooled to ∼ 45°C) was added to this mixture and immediately spread over BHI agar plate. Following overnight incubation at 37°C, plaques were further purified by repeating the plaque assay step (as mentioned above) two more times. Phage stock solutions were prepared by growing purified phages with their respective enterococcus hosts in 25 ml BHI broth in a shaker incubator maintained at 37°C overnight. Cultures were vortexed and then centrifuged at 10,000 × g for 10 minutes followed by supernatant filtration using 0.2 μm syringe filter. For long term storage, phages were stored at -80°C in BHI medium with 25% glycerol solution. To determine the phage titer, a plaque assay was performed with their respective experimental strains. Phage stocks were serially diluted up to 10^-8^ dilution using BHI broth and 5 μl from each dilution was spotted on BHI agar plates with an overlay of top agar containing host bacteria. Based on the plaque assay results, the titer of each phage (PFU/ml) was determined. Three of the total five phages used in this study (**Table 2**) were characterized previously by Wandro *et. al.* but the phage PL was isolated, purified, and characterized during this study.

### Phage and bacterial genome sequencing and analysis

Total genomic DNA from the phage PL and bacterial isolates were extracted using QIAamp UltraSens Virus kit (Qiagen catalog# 53706) and DNeasy Blood & Tissue Kit (Qiagen catalog# 69504) respectively. Quality of the extracted DNA was checked using Qubit dsDNA high sensitivity assay kit (Invitrogen, catalog# Q32851) and DNA libraries were prepared using Nextera XT DNA library preparation kit (Illumina, catalog# FC-131-1024). Paired end sequencing (2 × 150 bp) was used to sequence the whole genome of the phage PL on Illumina iSeq100 platform and the bacterial genomes were sequenced on Illumina Miseq platform. Sequencing reads were assembled into scaffolds using the DeNovo approach of CLC Genomics Workbench software version 21.0.3 (Qiagen, Redwood City, CA, USA). Next, genomes were annotated using an online open-source annotation tool RASTtk v2.0 (Rapid Annotation using Subsystem Technology tool kit [49]. Complete genome of the phage was visualized using PROKSEE analysis using CGView Server [50].

### Transmission Electron Microscopy

TEM staining and analysis was performed as previously described by Lee *et. al.,* with some modifications [51]. Briefly, 10 μl drops of concentrated phage solution (∼2 × 10^8^ PFU/ml) were spotted onto a clean parafilm sheet. Carbon-coated copper grids (PELCO SynapTek™ Grids, product# 01754-F) were placed over the drops for approximately one minute followed by three passes over the 20 μl drops of sterile deionized water and excess water was blotted using filter paper. Finally, the grids were negatively stained by placing over 10 μl drop of 2 % uranyl acetate solution (pH 4.0) for ∼ 45 seconds. The grids were then immediately blotted using filter papers and air dried at room temperature for 5 minutes. Grids were imaged after at least 24 h of staining using Joel 1400 plus at the University of California, San Diego - Cellular and Molecular Medicine Electron Microscopy Core facility (RRID:SCR_022039).

### Phage-Antibiotic Synergy testing

Phage-Antibiotic Synergy (PAS) testing was performed in BHI medium as previously described by Liu *et. al.,* with some modifications [52]. The experiment was setup in a 96 well plate in a total volume of 200 µl growth medium containing varying concentration of antibiotics and phages (**Figure 2**). The antibiotics and phages were serially diluted 10-fold from top to bottom and from right to left for antibiotics and phages, respectively. This created a concentration gradient for antibiotics only (1^st^ column), phages only (2^nd^ row from bottom) and antibiotic/phage (central wells). Similarly, bacterial growth control and no growth control were also set up in triplicate wells (**Figure 2**). Briefly, A single bacterial colony was inoculated in 2 ml BHI broth followed by overnight incubation in a shaker incubator maintained at 37°C and 200 rpm. The next morning, the culture was diluted in BHI broth (1:400) followed by a short incubation of ∼3 h to get exponentially growing bacterial cells. The culture was then adjusted to OD_600_ = 0.1 (∼ 1×10^8^ CFU/ml) by diluting in BHI broth and 20 µl of the diluted culture was added to each well of the 96 well plate containing 140 μl of BHI broth to yield a final concentration of ∼ 1×10^7^ CFU/ml in each well. Next, the phage stock solution was serially diluted 10-fold in BHI broth ranging from ∼1×10^8^ to 1×10^3^ PFU/ml and 20 µl from each dilution was added to each well of column B to column G, respectively. This resulted in a 10-fold reduction in phage concentration in each column compared to their respective diluted phage stock samples. To be specific, each well of column B had a final concentration of ∼1×10^7^ PFU/ml and each well of column G had a final concentration of ∼1×10^2^ PFU/ml. This established a range of multiplicity of infection (MOI, which is defined as the ratio of the numbers of phage particles to the numbers of the host cells) across the columns (B to G) from 1 to 1×10^−5^. This was followed by the addition of 20 μl of serially diluted antibiotic stock solutions to each well which resulted in a final concentration ranging from 128 to 0.5 µg/ml (VAN), 64 to 0.25 µg/ml (AMP), and 32 to 0.125 µg/ml (LZD) from row 2 to 10 (**Figure 2**). The plates were incubated in a VERSAmax Microplate Reader for 18 hours and OD600 was measured every 15 minutes after shaking for 3 seconds. Antibiotic stock solutions were prepared fresh in ultrapure sterile water, filter sterilized using 0.22 µm filters and stored at 4°C. The PAS experiment was performed in three biological replicates. The PAS was identified based on the percentage growth reduction of bacteria in each well which was determined using the following formula as previously described [52].

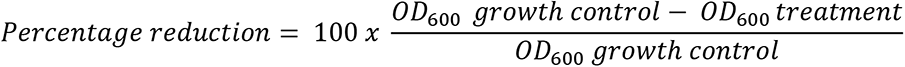

### Sampling scheme for RNA isolation and sequencing

To harvest bacterial samples for transcriptomic analysis, coculture assays were repeated in higher culture volume for specific growth conditions for *E. faecium* EF98PII and *E. faecalis* EF116PII which exhibited synergistic interaction during co-culture assays (see Fig. 3 and S1 for synergy conditions). Plates containing 24 wells (Thermo Scientific™ Nunc™ Non-Treated Multidishes, catalog# 144530) were used for setting up co-culture assay in a total volume of 600 μl BHI broth for each synergy combination along with proper controls for 5 hour and 18 hour time points. This experiment was repeated on three separate days that represented three biological replicates for each sample. Samples (16 samples × 3 biological replicates, n=48) were collected for each of the two bacterial strains which added to a total of 96 samples as shown in the sampling scheme (Fig. 4A). From each well, 450 μl sample was pooled into separate Eppendorf tubes and total RNA was extracted using the RNeasy Protect Mini Kit (Qiagen, catalog# 74124). Isolated RNA samples were quantified using both Qubit dsDNA HS (High Sensitivity) assay kit (Invitrogen, catalog# Q32851) and NanoDrop™ 2000/2000c (Thermo Fisher Scientific, catalog# ND2000CLAPTOP). Approximately 880 ng of RNA was used to build cDNA sequencing library using Illumina stranded total RNA prep with ribo zero plus kit (catalog# 20040529). 100 bp paired end sequencing was performed on Illumina NovaSeq 6000 platform using S4 flow cell (Illumina Inc., San Diego, CA, USA).

### Differential gene expression analysis

RNA sequencing analysis was performed using command line tools and R-studio (v2022.12.0+353). Initially, demultiplexed raw sequencing reads were trimmed using Trimmomatic (v0.39) with following parameters: ILLUMINACLIP:TruSeq3-PE.fa:2:30:10 LEADING:3 TRAILING:3 SLIDINGWINDOW:4:15 MINLEN:35 [53]. Reference genome file (FASTA format) and gene annotation files (GFF format) were downloaded from Nucleotide Sequence Database of NCBI for *Enterococcus faecium* NRRL B-2354 and EnsemblBacteria for *Enterococcus faecalis* V583. Reference genomes were indexed using HISAT2 (2.0.5) followed by sequencing read mapping [54]. Next, transcripts were assembled from mapped reads and reference transcript annotation files by StringTie [55] and transcript abundance in each sample was estimated and counted to identify differentially expressed genes using Ballgown tool [56]. Finally, DESeq2 algorithm of the free online server, iDEP.96 was used to calculate fold change values for differentially expressed genes [57]. A minimum value two-fold change was considered to filter out significantly upregulated and downregulated genes with an FDR cutoff value of less than 0.05. Finally, default parameters were used to perform pathway analysis using Gene Ontology (GO) database [58] and STRING: a database of predicted functional associations between proteins [59].

### Statistical analysis

All the experiments were performed in three independent biological replicates. The data shown in the heatmaps represents the mean percentage reduction of bacterial growth that originated from three biological replicates. Growth curve figures, and bar graphs show the means and standard deviations (SDs) of three biological replicates. To determine statistical significance one-way ANOVA was performed using Dunnett’s multiple comparisons test on GraphPad Prism 9 (v9.2.0) between treatment and control groups and a p-value of < 0.05 represented that the given data is significant.

### Data Access

Sequencing data uploaded to NCBI SRA: BioProject Accession# PRJNA947989 96 samples. RNA sequences of 96 bacterial samples (BioSample# SAMN33872066 - SAMN33872161), Whole genome sequences of 3 bacterial strains (BioSample# SAMN36748519, SAMN36748517, & SAMN36748518), Whole genome sequences of Phage PL (Biosample# SAMN36749450).

## References

1. Ventola CL. The antibiotic resistance crisis: part 1: causes and threats. P T. 2015;40(4):277–83.

2. CDC: Antibiotic resistance threats in the United States, 2019. In. Atlanta, GA: Centers for Disease Control and Prevention; 2019.

3. Bereket W, Hemalatha K, Getenet B, Wondwossen T, Solomon A, Zeynudin A, Kannan S. Update on bacterial nosocomial infections. Eur Rev Med Pharmacol Sci. 2012;16(8):1039–44.

4. Van Tyne D, Gilmore MS. Friend turned foe: evolution of enterococcal virulence and antibiotic resistance. Annu Rev Microbiol. 2014;68:337–56; doi: 10.1146/annurev-micro-091213-113003.

5. Cabiltes I, Coghill S, Bowe SJ, Athan E. Enterococcal bacteraemia ’silent but deadly’: a population-based cohort study. Intern Med J. 2020;50(4):434–40; doi: 10.1111/imj.14396.

6. Marothi YA, Agnihotri H, Dubey D. Enterococcal resistance--an overview. Indian J Med Microbiol. 2005;23(4):214–9.

7. Kristich CJ, Rice LB, Arias CA. Enterococcal Infection-Treatment and Antibiotic Resistance. In: Gilmore MS, Clewell DB, Ike Y, Shankar N, editors. Enterococci: From Commensals to Leading Causes of Drug Resistant Infection. Boston; 2014.

8. NI AH, Huycke MM. Enterococcal disease, epidemiology, and implications for treatment. 2014.

9. Arias CA, Murray BE. The rise of the Enterococcus: beyond vancomycin resistance. Nat Rev Microbiol. 2012;10(4):266–78; doi: 10.1038/nrmicro2761.

10. Perl TM. The threat of vancomycin resistance. Am J Med. 1999;106(5A):26S–37S; discussion 48S-52S; doi: 10.1016/s0002-9343(98)00354-4.

11. Hierholzer WJ, Garner JS, Adams AB, Craven DE, Fleming DW, Forlenza SW, et al. Recommendations for preventing the spread of vancomycin resistance: recommendations of the Hospital Infection Control Practices Advisory Committee (HICPAC). Am J Infect Control. 1995;23:87–94.

12. Weiner LM, Webb AK, Limbago B, Dudeck MA, Patel J, Kallen AJ, et al. Antimicrobial-Resistant Pathogens Associated With Healthcare-Associated Infections: Summary of Data Reported to the National Healthcare Safety Network at the Centers for Disease Control and Prevention, 2011-2014. Infect Control Hosp Epidemiol. 2016;37(11):1288–301; doi: 10.1017/ice.2016.174.

13. O’Driscoll T, Crank CW. Vancomycin-resistant enterococcal infections: epidemiology, clinical manifestations, and optimal management. Infection and drug resistance. 2015:217–30.

14. Parasion S, Kwiatek M, Gryko R, Mizak L, Malm A. Bacteriophages as an alternative strategy for fighting biofilm development. Pol J Microbiol. 2014;63(2):137–45.

15. Harper D, Parracho H, Walker J, Sharp R, Hughes G, Werthén M, et al. Bacteriophages and Biofilms. Antibiotics. 2014;3(3):270–84; doi: 10.3390/antibiotics3030270.

16. Kumaran D, Taha M, Yi Q, Ramirez-Arcos S, Diallo J-S, Carli A, Abdelbary H. Does treatment order matter? Investigating the ability of bacteriophage to augment antibiotic activity against Staphylococcus aureus biofilms. Frontiers in microbiology. 2018;9:127.

17. Morrisette T, Kebriaei R, Lev KL, Morales S, Rybak MJ. Bacteriophage Therapeutics: A Primer for Clinicians on Phage-Antibiotic Combinations. Pharmacotherapy. 2020;40(2):153–68; doi: 10.1002/phar.2358.

18. Khalifa L, Brosh Y, Gelman D, Coppenhagen-Glazer S, Beyth S, Poradosu-Cohen R, et al. Targeting Enterococcus faecalis biofilms with phage therapy. Appl Environ Microbiol. 2015;81(8):2696–705; doi: 10.1128/AEM.00096-15.

19. Chatterjee A, Johnson CN, Luong P, Hullahalli K, McBride SW, Schubert AM, et al. Bacteriophage Resistance Alters Antibiotic-Mediated Intestinal Expansion of Enterococci. Infect Immun. 2019;87(6); doi: 10.1128/IAI.00085-19.

20. Yoong P, Schuch R, Nelson D, Fischetti VA. Identification of a broadly active phage lytic enzyme with lethal activity against antibiotic-resistant Enterococcus faecalis and Enterococcus faecium. J Bacteriol. 2004;186(14):4808–12; doi: 10.1128/JB.186.14.4808-4812.2004.

21. Cheng M, Liang J, Zhang Y, Hu L, Gong P, Cai R, et al. The bacteriophage EF-P29 efficiently protects against lethal vancomycin-resistant Enterococcus faecalis and alleviates gut microbiota imbalance in a murine bacteremia model. Frontiers in microbiology. 2017;8:837.

22. Oliveira A, Sillankorva S, Quinta R, Henriques A, Sereno R, Azeredo J. Isolation and characterization of bacteriophages for avian pathogenic E. coli strains. Journal of applied microbiology. 2009;106(6):1919–27.

23. Capparelli R, Parlato M, Borriello G, Salvatore P, Iannelli D. Experimental phage therapy against Staphylococcus aureus in mice. Antimicrobial agents and chemotherapy. 2007;51(8):2765–73.

24. Torres-Barcelo C. Phage Therapy Faces Evolutionary Challenges. Viruses. 2018;10(6); doi: 10.3390/v10060323.

25. Oechslin F. Resistance Development to Bacteriophages Occurring during Bacteriophage Therapy. Viruses. 2018;10(7); doi: 10.3390/v10070351.

26. Lev K, Kunz Coyne AJ, Kebriaei R, Morrisette T, Stamper K, Holger DJ, et al. Evaluation of Bacteriophage-Antibiotic Combination Therapy for Biofilm-Embedded MDR Enterococcus faecium. Antibiotics. 2022;11(3):392.

27. Kebriaei R, Lev K, Morrisette T, Stamper KC, Abdul-Mutakabbir JC, Lehman SM, et al. Bacteriophage-antibiotic combination strategy: an alternative against methicillin-resistant phenotypes of Staphylococcus aureus. Antimicrobial Agents and Chemotherapy. 2020;64(7):e00461–20.

28. Lopatina A, Tal N, Sorek R. Abortive Infection: Bacterial Suicide as an Antiviral Immune Strategy. Annu Rev Virol. 2020;7(1):371–84; doi: 10.1146/annurev-virology-011620-040628.

29. Dy RL, Richter C, Salmond GP, Fineran PC. Remarkable Mechanisms in Microbes to Resist Phage Infections. Annu Rev Virol. 2014;1(1):307–31; doi: 10.1146/annurev-virology-031413-085500.

30. Khalifa L, Gelman D, Shlezinger M, Dessal AL, Coppenhagen-Glazer S, Beyth N, Hazan R. Defeating Antibiotic- and Phage-Resistant Enterococcus faecalis Using a Phage Cocktail in Vitro and in a Clot Model. Front Microbiol. 2018;9:326; doi: 10.3389/fmicb.2018.00326.

31. Chan BK, Abedon ST, Loc-Carrillo C. Phage cocktails and the future of phage therapy. Future Microbiol. 2013;8(6):769–83; doi: 10.2217/fmb.13.47.

32. Abedon ST, Danis-Wlodarczyk KM, Wozniak DJ. Phage Cocktail Development for Bacteriophage Therapy: Toward Improving Spectrum of Activity Breadth and Depth. Pharmaceuticals. 2021;14(10):1019; doi: 10.3390/ph14101019.

33. Wandro S, Ghatbale P, Attai H, Hendrickson C, Samillano C, Suh J, et al. Phage Cocktails Constrain the Growth of Enterococcus. mSystems. 2022;7(4):e0001922; doi: 10.1128/msystems.00019-22.

34. McCallin S, Brüssow H. Clinical Trials of Bacteriophage Therapeutics. In: Springer International Publishing; 2021. p. 1099-127.

35. Liu D, Van Belleghem JD, de Vries CR, Burgener E, Chen Q, Manasherob R, et al. The Safety and Toxicity of Phage Therapy: A Review of Animal and Clinical Studies. Viruses. 2021;13(7); doi: 10.3390/v13071268.

36. Lood C, Haas PJ, van Noort V, Lavigne R. Shopping for phages? Unpacking design rules for therapeutic phage cocktails. Curr Opin Virol. 2022;52:236–43; doi: 10.1016/j.coviro.2021.12.011.

37. Flores CO, Meyer JR, Valverde S, Farr L, Weitz JS. Statistical structure of host-phage interactions. Proc Natl Acad Sci U S A. 2011;108(28):E288–97; doi: 10.1073/pnas.1101595108.

38. Chan BK, Sistrom M, Wertz JE, Kortright KE, Narayan D, Turner PE. Phage selection restores antibiotic sensitivity in MDR Pseudomonas aeruginosa. Sci Rep. 2016;6:26717; doi: 10.1038/srep26717.

39. Tornheim JA, Dooley KE. The Global Landscape of Tuberculosis Therapeutics. Annu Rev Med. 2019;70:105–20; doi: 10.1146/annurev-med-040717-051150.

40. Gunthard HF, Saag MS, Benson CA, del Rio C, Eron JJ, Gallant JE, et al. Antiretroviral Drugs for Treatment and Prevention of HIV Infection in Adults: 2016 Recommendations of the International Antiviral Society-USA Panel. JAMA. 2016;316(2):191–210; doi: 10.1001/jama.2016.8900.

41. Horner C, Mushtaq S, Allen M, Hope R, Gerver S, Longshaw C, et al. Replacement of Enterococcus faecalis by Enterococcus faecium as the predominant enterococcus in UK bacteraemias. JAC Antimicrob Resist. 2021;3(4):dlab185; doi: 10.1093/jacamr/dlab185.

42. Szklarczyk D, Kirsch R, Koutrouli M, Nastou K, Mehryary F, Hachilif R, et al. The STRING database in 2023: protein-protein association networks and functional enrichment analyses for any sequenced genome of interest. Nucleic Acids Res. 2023;51(D1):D638–D46; doi: 10.1093/nar/gkac1000.

43. Kutter E. Phage host range and efficiency of plating. Methods Mol Biol. 2009;501:141–9; doi: 10.1007/978-1-60327-164-6_14.

44. Gelman D, Beyth S, Lerer V, Adler K, Poradosu-Cohen R, Coppenhagen-Glazer S, Hazan R. Combined bacteriophages and antibiotics as an efficient therapy against VRE Enterococcus faecalis in a mouse model. Res Microbiol. 2018;169(9):531–9; doi: 10.1016/j.resmic.2018.04.008.

45. Miller WR, Munita JM, Arias CA. Mechanisms of antibiotic resistance in enterococci. Expert Rev Anti Infect Ther. 2014;12(10):1221–36; doi: 10.1586/14787210.2014.956092.

46. Khong E, Oh J, Jimenez JM, Liu R, Dunham S, Monsibais A, et al. A simple solid media assay for detection of synergy between bacteriophages and antibiotics. bioRxiv. 2023; doi: 10.1101/2023.08.23.554535.

47. Selim S. Mechanisms of gram-positive vancomycin resistance (Review). Biomed Rep. 2022;16(1):7; doi: 10.3892/br.2021.1490.

48. Khan Mirzaei M, Nilsson AS. Isolation of phages for phage therapy: a comparison of spot tests and efficiency of plating analyses for determination of host range and efficacy. PLoS One. 2015;10(3):e0118557; doi: 10.1371/journal.pone.0118557.

49. Brettin T, Davis JJ, Disz T, Edwards RA, Gerdes S, Olsen GJ, et al. RASTtk: A modular and extensible implementation of the RAST algorithm for building custom annotation pipelines and annotating batches of genomes. Scientific Reports. 2015;5(1):8365; doi: 10.1038/srep08365.

50. Stothard P, Grant JR, Van Domselaar G. Visualizing and comparing circular genomes using the CGView family of tools. Briefings in Bioinformatics. 2019;20(4):1576–82.

51. Lee H, Ku H-J, Lee D-H, Kim Y-T, Shin H, Ryu S, Lee J-H. Characterization and genomic study of the novel bacteriophage HY01 infecting both Escherichia coli O157: H7 and Shigella flexneri: potential as a biocontrol agent in food. PloS one. 2016;11(12):e0168985.

52. Gu Liu C, Green SI, Min L, Clark JR, Salazar KC, Terwilliger AL, et al. Phage-antibiotic synergy is driven by a unique combination of antibacterial mechanism of action and stoichiometry. MBio. 2020;11(4):e01462–20.

53. Bolger AM, Lohse M, Usadel B. Trimmomatic: a flexible trimmer for Illumina sequence data. Bioinformatics. 2014;30(15):2114–20; doi: 10.1093/bioinformatics/btu170.

54. Kim D, Paggi JM, Park C, Bennett C, Salzberg SL. Graph-based genome alignment and genotyping with HISAT2 and HISAT-genotype. Nat Biotechnol. 2019;37(8):907–15; doi: 10.1038/s41587-019-0201-4.

55. Pertea M, Pertea GM, Antonescu CM, Chang TC, Mendell JT, Salzberg SL. StringTie enables improved reconstruction of a transcriptome from RNA-seq reads. Nat Biotechnol. 2015;33(3):290–5; doi: 10.1038/nbt.3122.

56. Frazee AC, Pertea G, Jaffe AE, Langmead B, Salzberg SL, Leek JT. Ballgown bridges the gap between transcriptome assembly and expression analysis. Nat Biotechnol. 2015;33(3):243–6; doi: 10.1038/nbt.3172.

57. Ge SX, Son EW, Yao R. iDEP: an integrated web application for differential expression and pathway analysis of RNA-Seq data. BMC Bioinformatics. 2018;19(1):534; doi: 10.1186/s12859-018-2486-6.

58. Ashburner M, Ball CA, Blake JA, Botstein D, Butler H, Cherry JM, et al. Gene ontology: tool for the unification of biology. Nature genetics. 2000;25(1):25–9.

59. Mering Cv, Huynen M, Jaeggi D, Schmidt S, Bork P, Snel B. STRING: a database of predicted functional associations between proteins. Nucleic Acids Research. 2003;31(1):258–61; doi: 10.1093/nar/gkg034.

